# Neurophysiological Features of STN LFP underlying Sleep Fragmentation in Parkinson’s Disease

**DOI:** 10.1101/2023.10.19.563075

**Authors:** Guokun Zhang, Huiling Yu, Yue Chen, Chen Gong, Hongwei Hao, Yi Guo, Shujun Xu, Yuhuan Zhang, Xuemei Yuan, Guoping Yin, Jianguo Zhang, Huiling Tan, Luming Li

## Abstract

**Background:** Sleep fragmentation is a persistent problem throughout the course of Parkinson’s disease (PD). However, the related neurophysiological patterns and the underlying mechanisms remained unclear.

**Method:** We recorded subthalamic nucleus (STN) local field potentials (LFPs) using DBS with real-time wireless recording capacity from thirteen PD patients undergoing a one-night polysomnography recording, one month after DBS surgery before initial programming and when the patients were Off-Medication. The STN LFP features that characterized different sleep stages, correlated with arousal and sleep fragmentation index, and preceded stage transitions during N2 and REM sleep were analyzed.

**Results:** Both beta and low gamma oscillations in NREM sleep increased with the severity of sleep disturbance (arousal index (ArI)-beta_NREM_: r=0.9,*p*=0.0001) and sleep fragmentation index (SFI)-beta_NREM_: r=0.6,*p*=0.0301; SFI-gamma_NREM_: r=0.6,*p*=0.0324). We next examined the low-to-high power ratio, which was the power ratio of theta oscillations to beta and low gamma oscillations, and found it to be an indicator of sleep fragmentation (ArI-LHPR_NREM_: r=-0.8,*p*=0.0053; ArI-LHPR_REM_: r=-0.6,*p*=0.0373; SFI-LHPR_NREM_: r=-0.7,*p*=0.0204; SFI-LHPR_REM_: r=-0.6,*p*=0.0428). In addition, long beta bursts (>0.25s) during NREM stage 2 were found preceding the completion of transition to stages with more cortical activities (towards Wake/N1/REM compared with towards N3 (*p*<0.01)) and negatively correlated with STN spindles, which were detected in STN LFPs with peak frequency distinguishable from long beta bursts (STN spindle: 11.5Hz, STN long beta bursts: 23.8Hz), in occupation during NREM sleep (β = -0.24,*p*<0.001).

**Conclusion:** Features of STN LFPs help explain neurophysiological mechanisms underlying sleep fragmentations in PD, which can inform new intervention for sleep dysfunction.

**What is already known on this topic:** Beta oscillation, which is a biomarker for rigidity and bradykinesia during awake, significantly reduced during NREM sleep compared to REM or awake. Researches from MPTP non-human primate model suggested increased beta oscillation in basal ganglia contributed to insomnia in PD, however evidence in human patients is still lacking.

**What this study adds:** Beta and low gamma band activities in STN LFPs during sleep recorded from human PD patients correlated with severity of sleep impairment. The low-high power ratio can serve as a biomarker for sleep fragmentation. Besides, pathological beta bursts and physiological sleep spindles can be detected in STN LFP during NREM sleep and distinguishable from each other.

**How this study might affect research, practice or policy:** These findings enhance our understanding of the electrophysiological mechanisms that underlie sleep fragmentation in Parkinson’s disease. The study also has implications on the design of closed-loop DBS: we may need to take multiple frequency band activities into consideration and differentiate pathological beta oscillation from the sleep spindles.

## 1. Introduction

Parkinson’s disease (PD) is one of the most prevalent neurodegenerative diseases predominately affecting dopaminergic system in basal ganglia (BG). Apart from prominent movement disorders such as bradykinesia, rigidity, tremor and balance disturbance, more than 80% PD patients suffer from sleep disturbance, which persists the entire disease course and worsens disease prognosis ^1,2^. Sleep fragmentations, in particular, correlate with sleep-maintenance insomnia and excessive daytime sleepiness, can further interact with cognitive impairment and mental illness, and lead to disease progression acceleration^3–6^. However, the mechanism of sleep fragmentation and underlying neuronal activities in the BG remain largely unknown and result in the obstacles towards developing sleep-specific intervention.

A recent study based on non-human primate model of Parkinson’s disease suggested that beta oscillations (13-30Hz) in the basal ganglia increased during NREM sleep in Parkinsonian state, which was associated with sleep fragmentation and reduction in slow wave activities^7^. However, evidence on the role of increased beta oscillation in insomnia is still missing in human patients, as most existing studies with LFP recordings from human patients show that the beta oscillation was significantly reduced during NREM sleep compared to REM or awake ^8,9^. Moreover, as wide spread sleep spindles, which were transient waxing and waning 11-16 Hz oscillations prominent during NREM stage 2 sleep, were found in different basal ganglia nucleus in healthy non-human primate^10^. Whether they are dissociable or are interfered by pathological beta bursts, the frequency of which were overlapped with spindles, remained unclear. Thus, gaining more knowledge on this topic will improve our understanding of the mechanism.

In the past decade, the appearance of DBS with recording function acts an important technique for detecting the long-term changes of deep brain signals^11^, which can simultaneously collect local field potentials (LFP) around electrodes while stimulate specific brain regions^12^. In our previous studies, we have reported a novel design of DBS system combining DBS therapy as well as concurrent measurement and real-time LFP transmission together^13,14^. Through in vitro wireless recharging technology, the system also broke through the limitation of acquisition time, established a long-term, safe and stable detection of human deep brain technology^15,16^. Here, based on this technology, we conducted simultaneous polysomnography monitoring together with STN LFP recording during a whole-night sleep in 13 patients with Parkinson’s disease to investigate the neurophysiology mechanism of STN LFPs underlying sleep fragmentation in PD.

## 2. Methods

### Participants and Clinical Evaluation

The protocol of this study was approved by the ethical committees of the surgical hospitals (Beijing Tiantan Hospital of Capital Medical University, Peking Union Medical College Hospital, and Qilu Hospital of Shandong University) and registered at ClinicalTrials.gov (Identifier NCT02937727). A total of 13 patients (8 males) with PD with the average age of 56±6.7 years old and average disease duration of 9.5±4.3 years were recruited in this study. All patients had signed informed consent in line with the Declaration of the Principles of Helsinki prior to surgery. All patients were bilaterally implanted in STN with electrodes consisted of four contacts (Model L301C, Beijing PINS Medical Co, China) and a sensing-enabled neurostimulator (G106R, Beijing PINS Medical Co, China). Locations of the electrodes were identified based on presurgical structural MRI and postsurgical computed tomography (CT) and reconstructed with Lead DBS tool^17^ as shown in Figure 1a.

**Figure 1.**
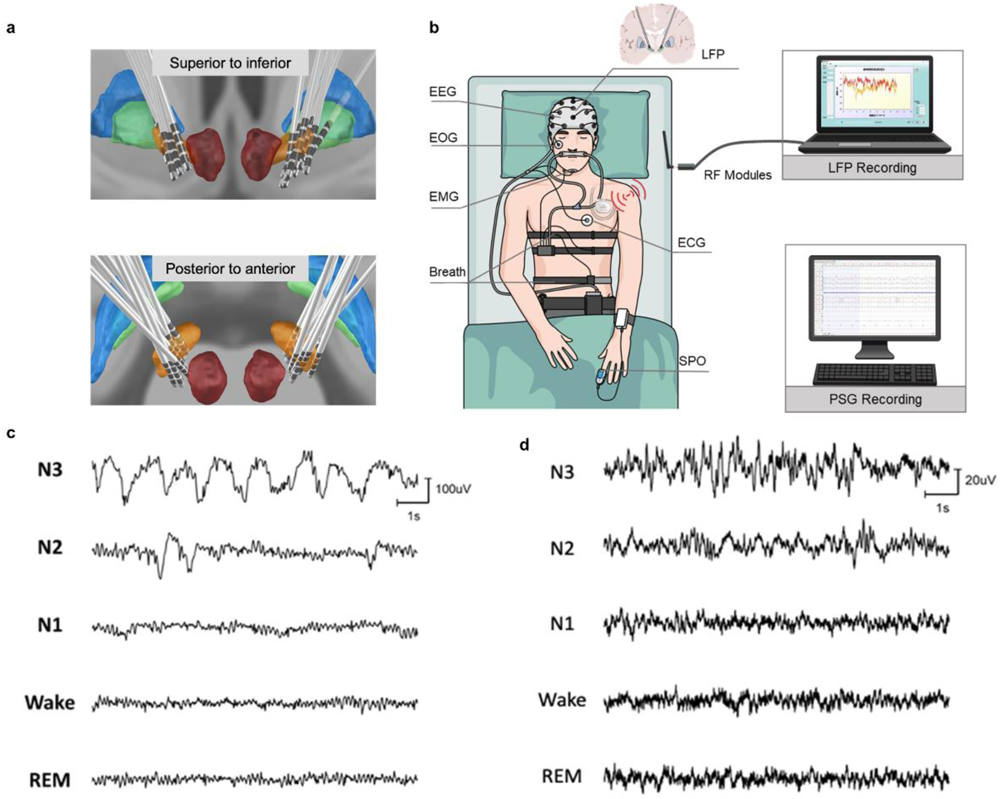
Lead Localization, Schematic of Recording Platform and Waveform Examples of EEG and STN LFP. a. Locations of electrodes from all 12 patients, for whom the LFPs were analysed, were reconstructed by Lead DBS and viewed from superior to inferior and posterior to anterior; b. The polysomnography system consisted of 6 channels of electroencephalogram (EEG), submental electromyogram (EMG), electrooculogram (EOG), and electrocardiogram (ECG) recordings. LFPs were recorded through the chronically implanted sensible DBS system and transmitted wirelessly to a PC with RF modules; c-d. Example waveforms of 5 sleep stages of EEG (c) and STN LFP signal (d) obtained from a single subject.

Clinical evaluation including Unified PD Rating Scale motor score (UPDRS III) Hoehn and Yahr Scale and Montreal Cognitive Assessment were conducted. One participant (subject 8) was excluded from the STN LFP analysis due to abnormal impedance of the contacts (>100KΩ), but was included in the analysis of sleep parameters. Demographic characteristics and clinical details and the patients included in the analysis are summarized in **Table 1**.

**Table 1.** Demographics of Participants.

| No. | Gender | Age | Disease duration | H-Y<sup>a</sup> | Estimated presurgery LEDDs | UPDRS III (off medication)<sup>b</sup> |  |  | MMSE<sup>d</sup> |  | Tremor (DBS off) | Record contacts | Treatment contacts |
| --- | --- | --- | --- | --- | --- | --- | --- | --- | --- | --- | --- | --- | --- |
|  |  |  |  |  |  | Pre-operation | DBS off<sup>c</sup> | DBS on | Pre-operation | 1 month |  |  |  |
| 1 | M | 53 | 13 | 3 | 189 | 34 | 41 | 11.5 | 25 | 28 | 0/0 | 2+4-/7+8-* | C+3-/C+5- |
| 2 | M | 67 | 8 | 3 | 800 | 20.5 | 16.5 | 2.5 | 27 | 26 | 0/0 | 2+4-/6+8- | C+3-/C+7- |
| 3 | M | 51 | 20 | 4 | 469 | 20 | 51 | 15.5 | 27 | 27 | 2/2 | 2+4-/5+7-* | C+3-/C+6- |
| 4 | M | 65 | 6 | 3 | 649 | 21 | 26 | 11.5 | 28 |  | 1/3 | 2+4-/5+7- | C+3-/7+8- |
| 5 | F | 60 | 7 | 3 | 800 | 28.5 | 20.5 | 9 | 28 | 27 | 0/0 | 2+4-/6+8-* | C+4-/C+7- |
| 6 | M | 46 | 7 | 4 | 201 | 46.5 | 36 | 17.5 | 29 |  | 0/0 | 1+3-/5+7-* | C+2-/C+6- |
| 7 | F | 61 | 8 | 4 | 620 | 29 | 33 | 15 | 22 | 24 | 2/0 | 2+4-/5+7- | C+3-/C+8- |
| 8<sup>e</sup> | M | 59 | 6 | 10 | 200 | 8 |  |  | 28 |  | 0/0 |  |  |
| 9 | F | 51 | 6 | 1.5 | 500 | 7.5 | 5.5 | 2 | 28 | 29 | 0/5 | 1+3-/7+8- | 4+3-/ C+8- |
| 10 | F | 47 | 12 | 5 | 810 | 38 | 37 | 18 | 28 | 29 | 2/2 | 2+4-/7+8- | C+3-/ C+5- |
| 11 | M | 56 | 15 | 4 | 300 | 50.5 | 40.5 | 18 | 26 | 28 | 2/2 | 1+3-/5+7-* | C+2-/ C+6- |
| 12 | M | 61 | 8 | 4 | 326 | 41 | 41.5 | 24 | 30 | 29 | 1/1 | 1+3-/5+7-* | C+2-/ C+6- |
| 13 | F | 51 | 8 | 4 | 443 | 53.5 | 51.5 | 12.5 |  | 26 | 0/0 | 2+4-/5+7-+ | C+3-/ C+6- |

2 H-Y<sup>a</sup>: Hoehn and Yahr Scale.
3 UPDRS III (off medication)<sup>b</sup>: The score was average of the results assessed by two movement disorders specialists according to the Unified Parkinson's Disease Rating Scale III (UPDRS III), and the difference of scores from the 4 two specialists was less than 10%.
5 DBS off<sup>c</sup>: The evaluations were conducted before initial programming.
6 MMSE<sup>d</sup>: Minimum Mental State Examination
7 8<sup>e</sup>: Subject 8 was excluded from the STN LFP analysis due to abnormal impedance of the contacts (>100KΩ), but was included in the analysis of sleep parameters
\*: Periodic oscillatory activities in spindle frequency band was identified
9 +: Periodic oscillatory activities in beta frequency band was identified

### Recording

Experiments were conducted one month after the DBS surgery before initial programming in a fully equipped sleep lab at the hospital. Patients were in Off-DBS and Off-Medication state, with long-acting antiparkinsonian medicine withdrawn for at least 24 hours and short-acting antiparkinsonian medicine withdrawn for at least 12 hours before the experiment. None of the patients involved in this study took drugs for sleep during the period of our research.

Together with STN LFP recording, Subjects underwent a whole-night polysomnography (PSG) monitoring which included 6 channels of EEG (F3-M2, F4-M1, C3-M2, C4-M1, O1-M2, O2-M1 according to the international 10-20 system), submental EMG, EOG and ECG recordings (Figure 1b). Signals of STN LFP were recorded wirelessly using a sensible DBS system and dedicated software (G106R, Beijing PINS Medical Co, China). The system was integrated the recording and real-time transmission technology with stimulation function^13,15^. In order to inform better control algorithms for closed-loop DBS to deal with sleep fragmentation in PD, bipolar signal adjacent to the stimulation contact, which is the commonly used feedback signal of closed-loop DBS, was used for the following analysis. LFP signals were sampled at 500Hz after filtered with an RC filter of 0.3Hz∼250Hz implemented on board. The distance of wireless transmission was up to 2m, avoiding the disturbance of normal sleep caused by LFP recording platform. The LFP recordings were synchronized with the PSG recordings through manually noting the recording starting time of each recording systems. Therefore, the resolution of the synchronization between the two recording systems is about 1 second.

### Sleep Evaluation

The polysomnography recording was evaluated by two sleep experts, who independently labeled each 30s-epoch data as Wake, REM sleep, N1, N2 or N3 according to the manual guideline from the American Academy of Sleep Medicine (AASM). Only the epochs that had the same labelling from the two experts were considered in this study. The validated sleep stages were also further served as reference for LFP signal analysis after correcting systemic time errors of PSG and LFP recording platforms. Measurement of sleep fragmentation and analysis of EMG and EOG were listed in Supplementary Methods.

### Data Processing

For EEG signals, C3-M2 and C4-M1 channels were selected for analysis after resampled at 256Hz. For LFP signals, to minimize the effect of volume conduction, we constructed a bipolar LFP signal for each hemisphere based on recordings from pairs of contacts adjacent to the contact used for therapeutic stimulation. In cases where the stimulation contact was at the end of the electrode (either contact 0 or 3, See Table 1), the bipolar channel that showed highest beta peak in the PSD when the patients were at rest and awake, were used for further analyses.

EEG and LFP signals were preprocessed to reduce movement artifacts and 50Hz power line interference (See Supplementary Methods). Moving window Short-time Fourier transform was used to estimate power spectrum density (PSD) of the EEG and LFP signals of each time point, with 2 second hamming window, 95% overlap, and a frequency resolution of 1Hz. Average power of typical oscillation bands (delta, 1-3Hz; theta, 4-7Hz; alpha 8-14Hz; beta, 15-30Hz; low gamma, 31-60Hz) were then extracted^27^.

### Spindle and Beta Burst Determination

In order to investigate the relationship between the beta bursts and sleep spindles during NREM sleep, in this study, both spindle and beta bursts detections were only performed on hemispheres with prominent beta (10-35Hz) or spindle activity (10-17Hz) during Wake and NREM, specifically, identified using FOOOF algorithm^18^ (more details in the Supplementary Material). Periodic oscillatory activities in spindle and beta frequency bands were identified in 11 and 18 out of the 24 recorded hemispheres, respectively. This resulted in eleven out of the 24 hemispheres (from nine patients) showed both distinct peak in the spindle frequency band during NREM sleep and the beta frequency band during awake.

Spindle events detection was performed with an algorithm which was widely applied and verified in previous studies of LFP and EEG^10,19^. Beta bursts were identified as previous illustrated^20^. Only beta bursts longer than 250ms were included in our analysis. See more details in Supplementary Methods.

### Statistical Analysis

We employed Wilcoxon rank-sum test to compare the electrophysiological activities between different sleep stages. FDR correction were performed in multiple comparisons. Correlation analysis in this study were conducted by Spearman’s rank correlation coefficient. Linear mixed-effects models (LMM) were used to compare the differences in electrophysiological characteristics. In each comparison, recorded hemispheres were considered as random effects to compensate multiple measurements within and between hemispheres. All data processing and statistical analysis were performed in MATLAB 2021b.

## 3. Results

### 3.1 Abnormal Sleep Architectures Correlated with Clinical Impairment in PD Patients

The percentage of different sleep stage time during the whole-night sleep was analyzed according to PSG evaluation. Compared with health control within the similar age group as shown in previous research, PD patients showed a tendency of increased percentage in Wake and NREM Stage 1 (N1) and decreased percentage in NREM Stage 3 (N3). Across recorded participants, the sleep efficiency of this recorded night correlated with cognitive function, as measured by the score of MMSE tested the day before PSG monitoring (N = 10, r=0.7, *p*=0.0392, Figure 2c), but not with UPDRS III (N = 12, r=0.2, p>0.1, Figure 2d). Correlation analysis showed that N3 percentage and N3 transition patterns, including the N2-to-N3 transition probability and N3 stability, all negatively correlated with UPDRS III score (N = 12, r=-0.6, *p*=0.0396, Figure 2e for N3 percentage; r=-0.6, p=0.0291, Figure2f for N2-to-N3 transition probability; and r=-0.6, *p*=0.0516, Figure 2g for N3 stability), indicating the essential association between motor impairment and shortened and fragmented N3 sleep.

**Figure 2.**
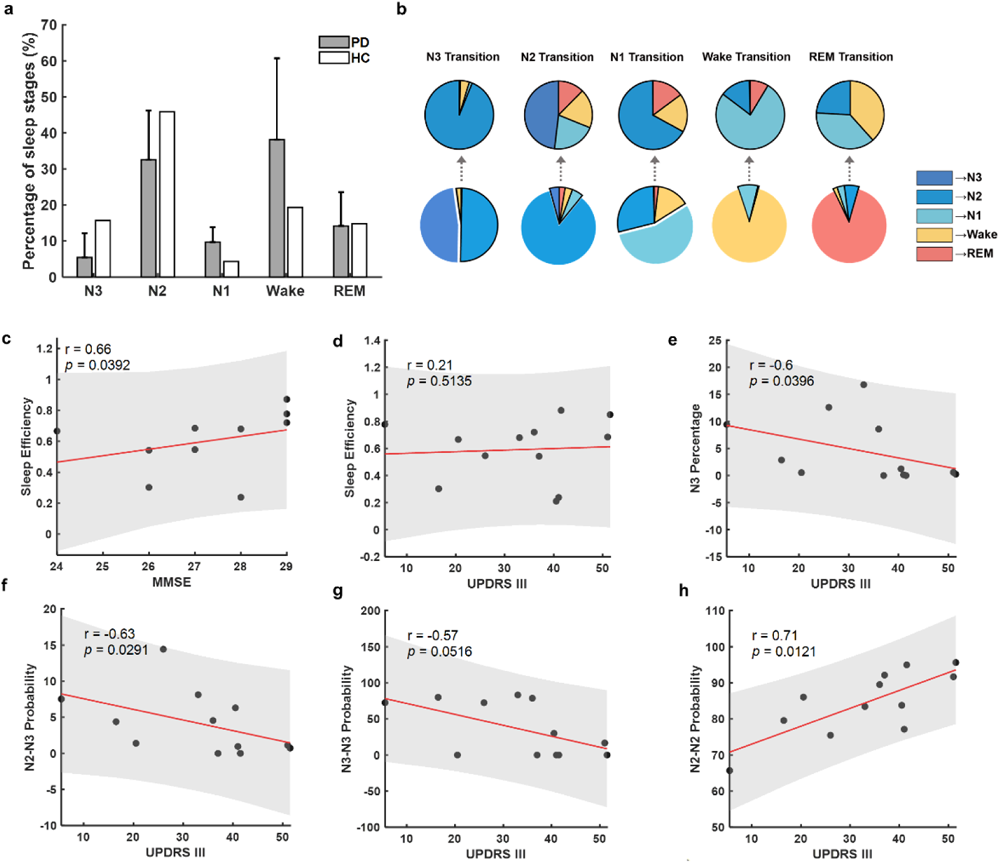
Sleep Parameters and Clinical Correlation a. Comparison of sleep stage percentage between Parkinson patients (grey bars) and health control (white bars) within the similar age group. The grey bar illustrated the mean± SEM of sleep stage percentage from 13 patients; b. Average sleep stage transition probabilities across sleep stages. The pie charts in the first raw presented the probabilities of sleep transition from one sleep stage to another different sleep stages; the second raw presented the probabilities of each sleep stage transitioning to another sleep stage in the following epoch. c. Correlations between sleep efficiency and MMSE. d. Correlations between sleep efficiency and UPDRS III. e. Correlations between N3 percentage and UPDRS III. f. Correlations between N2-N3 Transition Probability and UPDRS III. g. Correlations between N3-N3 Transition Probability and UPDRS III. h. Correlations between N2-N2 Transition Probability and UPDRS III. In c-h, each dot represents data from one participant; the red solid line and grey shading indicate the linear fitting and 95% confidence intervals.

### 3.2 Sleep-stage Dependent Characteristics of STN LFP

Through simultaneous PSG and STN LFP recording, characteristics of STN LFPs in different sleep stages were investigated. Average power spectrum density (PSD) of the central EEG and STN LFPs for different sleep stages were shown in Figure 3a and 3b, respectively. In STN LFP, frequency band of delta, theta and alpha showed elevated power with increasing sleep depth from Wake to NREM stages, and were highest during deep sleep stages (N2 and N3). In contrast, activities in beta and gamma bands presented an opposite pattern, which were highest during Wake compared with other sleep stages (Figure 3d). In REM sleep, the power of beta and gamma frequency bands were elevated compared with N2 and N3 stage, while no significant difference were shown between N1 and REM in the power of any of the frequency bands. In addition, no statistical significance was observed in the frequency band power of REM and Wake comparison, except for theta band, which were slightly higher in REM than in Wake stage. The results of statistical analysis were listed in Supplementary Table 3.

**Figure 3.**
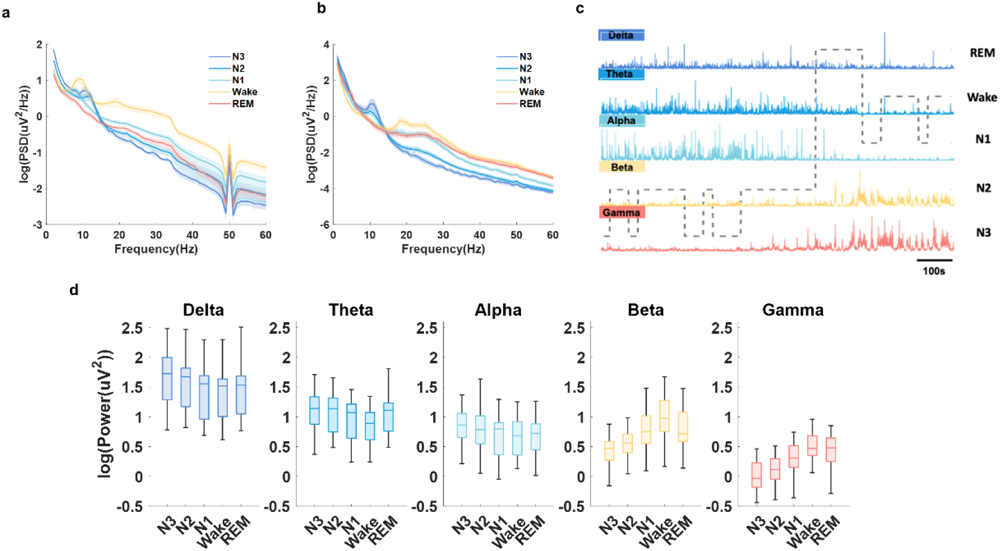
Sleep-stage Dependent Characteristics of STN LFP and EEG. a. Averaged PSDs (mean ± SEM) of different sleep stages from C3 and C4 channels of EEG; b. Averaged PSDs (mean ± SEM) of different sleep stages from LFP. The PSD results were averaged for all hemispheres; c. Illustration of how delta, theta, alpha, beta and gamma oscillations in STN LFPs changes with time and across different sleep stages in one exemplar patient; d. Box-and-whisker plots depicting changes in the power of delta, theta, alpha, beta, and gamma frequencies during different sleep stages The box edges represented the first quartile to the third quartile, with a vertical line drawing through the box at the median.

### 3.3 Increased subthalamic beta and Low gamma band activities contributed to Sleep Fragmentation

According to previous research, disrupted sleep in MPTP-induced nonhuman primate models of Parkinsonism was partially attributed to pathological neural activities, especially elevated beta oscillations, in basal ganglia. We next verify whether the phenomenon still exist in human STN. We used arousal index (Arl) and sleep fragmentation index (SFI) to quantify the severity of sleep fragmentation and disturbance. The detailed definition and calculation were listed in Supplementary Methods. In order to remove the confounding factor that the average power of different frequency bands was modulated by sleep stage, we considered the average power of different frequency bands for REM and NREM separately. The analyses showed that ArI and SFI correlated positively with average power of beta oscillation in NREM sleep (ArI-Beta_NREM_: r=0.9, *p*=0.0001;SFI-Beta_NREM_: r=0.64, *p*=0.0301;Figure 4a, d). The same trend was also found in low gamma oscillations (SFI-Gamma_NREM_: r=0.6, *p*=0.0324; Figure 4e). On the contrary, negative association between average theta oscillation power in NREM sleep and ArI (ArI-Theta_NREM_: r=-0.8, *p*=0.0047; Figure 4c). However, in REM sleep, only average power of low gamma oscillations was found to be positively correlated with SFI (SFI-Gamma_REM_: r=0.6, *p*=0.0428; Supplementary Figure 2e).

**Figure 4.**
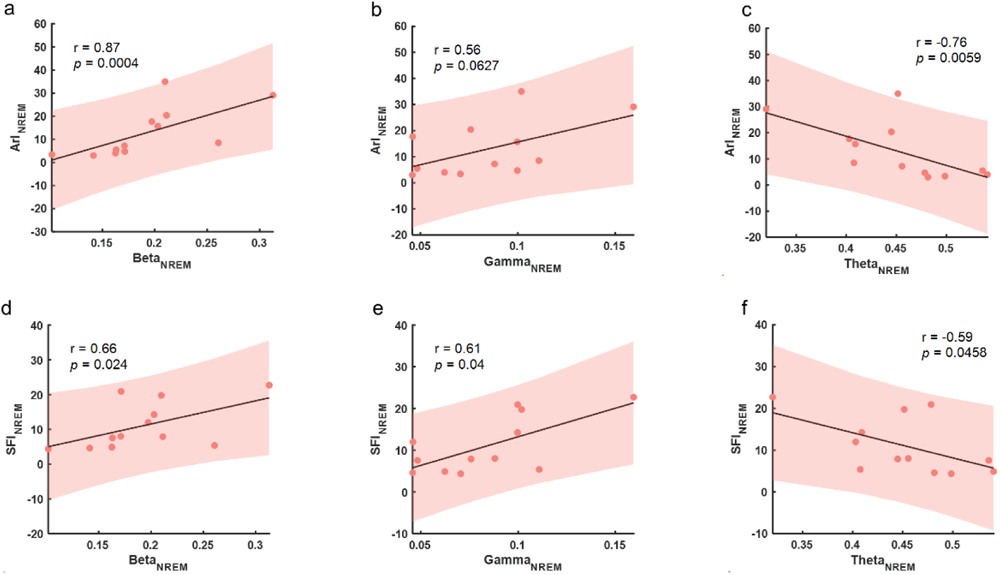
Correlation between Sleep Fragmentation Severity and STN LFP Oscillations. a-c. Correlations between arousal index and average power of beta, gamma and theta oscillations during NREM sleep. d-e. Correlations between sleep fragmentation index and average power of beta, gamma and theta oscillations during NREM sleep. In a-f, each dot represents data from one participant (the LFP features were averaged across the two hemispheres for each participant); the grey solid line and red shading indicate the linear fittings and 95% Confidence intervals were shown.

We further investigated how beta and low gamma band activities participated in sleep fragmentation. As sleep is a dynamic process, we next investigated the neurophysiological correlates of the transition processes during N2 and REM stages. In particular, we focused on the 120s of data during N2 and REM just before switching to other sleep stages. We categorized the sleep transition process from N2 to N3 (N2-N3), N1 (N2-N1), Wake (N2-Wake), N2 (N2-REM), as well as REM to N2 (REM-N2), N1 (REM-N1), Wake (REM-Wake). The sample size of events in each transition condition are listed in Supplementary Table 3. Low-to-high power ratio (LHPR) of STN LFPs, which is the ratio of the total power of low frequency oscillations (theta) divided by the total power of high frequency oscillations (beta and gamma), were analyzed and compared for each type of transition. In both N2 and REM transition analysis, statistical significance was shown in each of the non-overlapping 30s epochs before the transition completed. The LHPR were higher preceding the transition into deeper sleep stages, as during N2-N3 and REM-N2 process, compared to the similar time window preceding the transition to lighter sleep stages (Figure 5a, b). To explore the changes with more refined time resolution, LHPR at each time step (with 10 s moving window for each calculation and 0.1 second step length) preceding the sleep stage transition was plotted. As shown in Figure 5c, the LHPR gradually reduced with time before both N2-REM, N2-N1 and N2-Wake transitions, but kept high in N2-N3 transitions (Figure 5c). In REM transition, the LHPR remained higher in REM-N2 transition comparing with REM-N1 and REM-Wake transitions (Figure 5d). Overall, the average LHPR in both NREM and REM sleep negatively correlated with ArI (LHPR_NREM_: r=-0.8, *p*=0.0053; LHPR_REM_: r=-0.6, *p*=0.0373) and SFI (LHPR_NREM_: r=-0.7, *p*=0.0204; LHPR_REM_: r=-0.6, *p*=0.0428) (Figure 5 e-h), suggesting that the lower LHPR in STN LFP during sleep, the higher the sleep fragmentation in PD.

**Figure 5.**
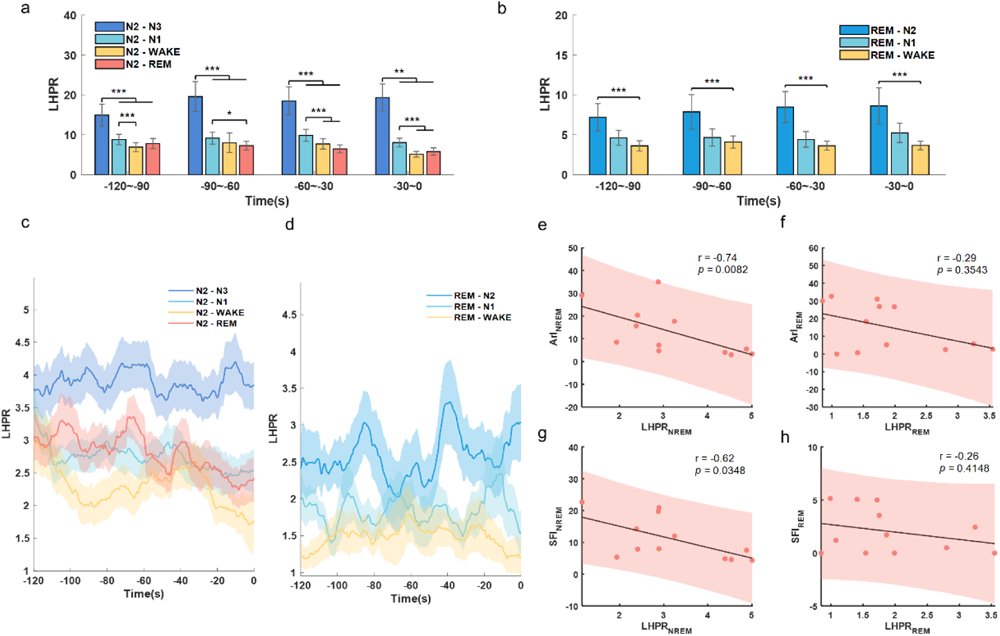
Analysis of LHPR during N2 and REM Transition Process. a. Comparison of LHPR preceding N2-N3, N2-N1, N2-Wake, and N2-REM transitions. b. Comparison of LHPR preceding REM-N2, REM-N1 and REM-Wake transitions. In both panel a and panel b, the bar graphs illustrated the mean± SEM of LHPR for all hemispheres in non-overlapping 30-second epochs, from 120 seconds to 0 seconds before completion of the transitions. **, *p*<0.01; ***, *p*<0.001; c-d. LHPR changes with time before different transitions from N2 and REM, respectively. In c-d, time 0 indicates the change of sleep stage label, the solid lines and shades represent the mean± SEM across hemispheres. e-f. Correlations between sleep fragmentation index/arousal index and the average LHPR of NREM sleep. g-h. Correlations between sleep fragmentation index/arousal index and average LHPR of REM sleep. In e-h, each dot represents data from one participant (LFP features were averaged across the two hemispheres for each participant); the grey solid line and red shading indicate the linear fittings and 95% Confidence intervals were shown.

### 3.4 Beta burst preceded transition to sleep stages with more cortical activity and interfered with sleep spindles

Long beta burst (>0.25s, briefly referred as beta burst) has been found to be a pathological biomarker of bradykinesia and rigidity in PD. However, whether it also participated in sleep disturbance in PD remains unknown. Here we found beta burst can still be detected during NREM, even though the density and occupation of beta bursts in N1, Wake and REM stages were significantly higher than N2 and N3 stages (*p*<0.05) (Figure 6 a-c). Beta bursts during different transitions in N2 and REM were then analyzed. During the last 120 seconds preceding the completion of N2 transition, the occupations of beta bursts were higher before transiting to stages with more cortical activity (N2-Wake, N2-N1 and N2-REM) than preceding the transition to deeper sleep stage (N2-N3 transitions) (*p*<0.01; Figure 6d). No difference in beta bursts were observed for different REM transitions (Figure 6e).

**Figure 6.**
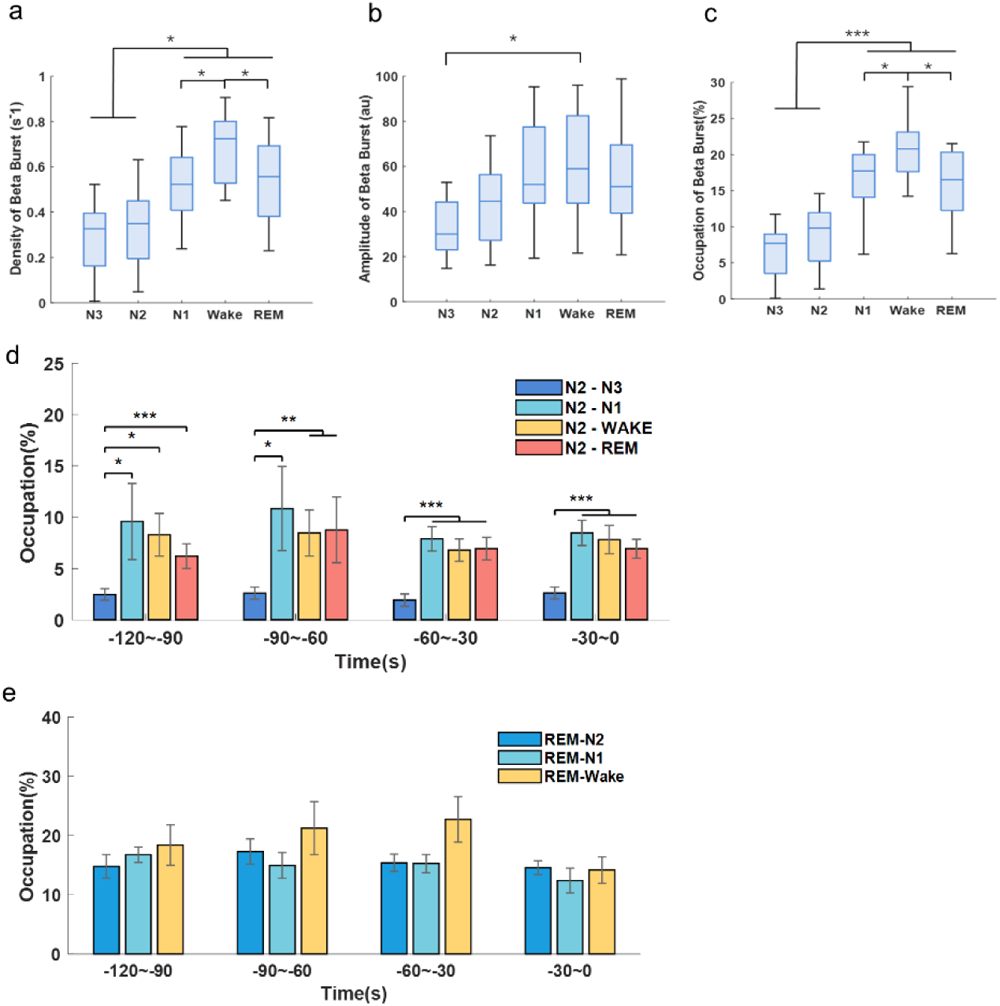
Long Beta Burst of STN LFP during Sleep. a-c. The density (number of events per second), duration and occupations of long Beta bursts (>0.25s) during different sleep stages. The box edges represent the first quartile to the third quartile, with a vertical line drawn through the box showing the median of all hemispheres. *, *p*<0.05; ***, *p*<0.001; d. Comparison of long Beta bursts occupations between N2-N3, N2-N1, N2-Wake, and N2-REM transitions. e. Comparison of long Beta bursts occupations between REM-N2, REM-N1, REM-Wake transitions. In d and e, the bar graphs illustrate the mean± SEM of long Beta bursts occupations for all hemispheres in non-overlapping 30-second epochs, from 120 seconds to 0 seconds before completion of the transitions. No statistical significance was shown.

As we observed both beta bursts and sleep spindles in NREM sleep, we explored how they interacted with each other. We compared the central frequency of beta burst and spindles in both cortex and STN, and found a clear distinction in the peak frequency ranges among beta bursts (23.8Hz) and spindles in both STN (11.5 Hz) and cortex (C3-M1 channel, 12.7 Hz) (Figure 7e). In addition, average occupations (in percentage of time) of cortical and subthalamic spindles as well as beta bursts were calculated over each 120s window. Linear mixed-effects models (LMM) identified negative correlations between the occupation of beta bursts and spindles in both STN (β = -0.24, *p*<0.001) and cortex (β = -0.15, *p*<0.001) (Figure 7f,g).

**Figure 7.**
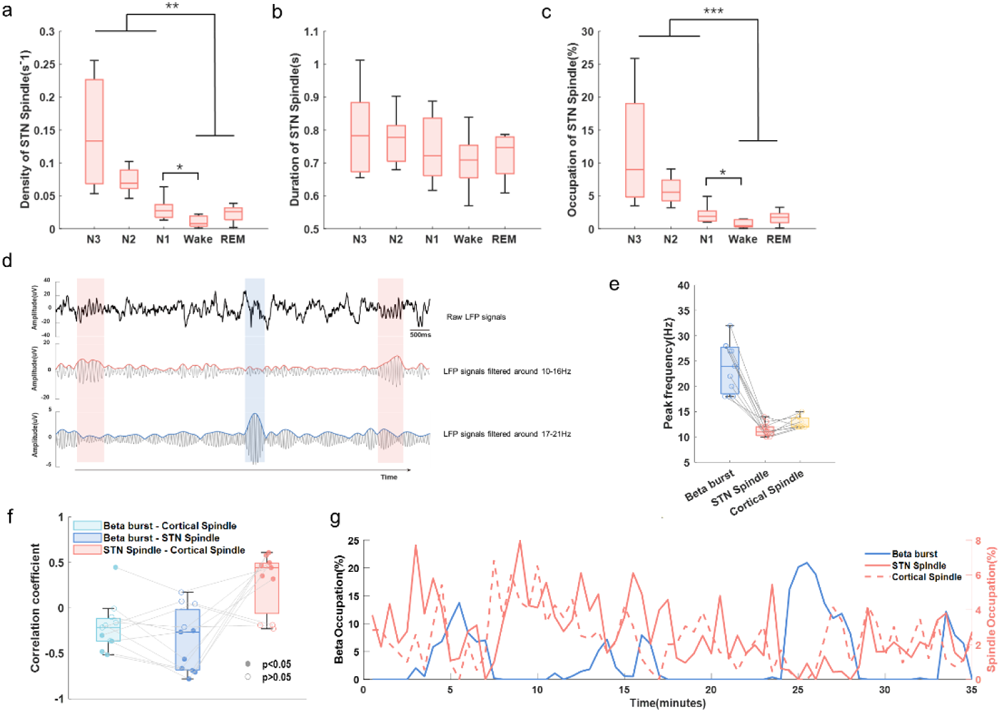
Interaction between Long Beta Burst and Sleep Spindles. a-c. The density, duration and occupations of sleep spindles of STN LFP in different sleep stages. The box edges represent the first quartile to the third quartile, with a vertical line drawn through the box showing the median of all hemispheres. **, *p*<0.01; ***, *p*<0.001; d. Example of detected sleep spindles and long Beta bursts of STN LFP during NREM sleep in one hemisphere. The detected beta burst and spindles were marked by blue and red shadow respectively. These three subplots showed the raw LFP signal (top), the 10-16Hz bandpass filtered signal and its envelope (middle), and the 17-21Hz bandpass filtered signal and its envelope in sequence (bottom); e. The peak frequency of beta and sleep spindles in both STN and cortex were distinguishable from each other; f. The Spearman correlation coefficients between the occupation of STN Beta burst and cortical spindle, STN Beta bursts and STN spindle, as well as STN spindles and cortical spindles, respectively for hemispheres in which both prominent beta and spindles were detected. The filled dots represent the coefficients with a *p* value less than 0.05 while the unfilled dots represent coefficients with a *p* value greater than 0.05; g. A typical example of the occupation of long beta bursts, STN spindles and cortical spindles changes over time during NREM sleep from a single subject. The occupations of spindles and beta bursts were averaged over each 120s window and concatenated during NREM sleep.

## 4. Discussion

In this study, we characterized neurophysiological features of STN LFPs underlying sleep fragmentation and abnormality. These results offer new insights into the mechanisms of sleep fragmentations in PD and provide guidance on further interventions.

### Features in STN LFPs underlying sleep fragmentation and insomnia

How neural activities in the subcortical structures contribute to sleep disorders remains largely unknown. Here we found that the average power of both beta and low gamma oscillations in STN LFPs during NREM sleep were positively correlated with the severity of sleep fragmentation. This is consistent with the behavioral correlation between the motor impairment (UPDRS III scores) and N3 sleep duration and continuity, suggesting that sleep fragmentation in PD can be related to motor impairments and its neural biomarker.. These results were also partially consistent with the previous observations from MPTP-primate models that increased beta oscillations in the STN LFP was associated with sleep fragmentation and decreased NREM sleep. Besides, as theta oscillations in STN LFP have been reported to be positively associated with cognitive functions, interruption in theta oscillation could be another mechanism underlying sleep disturbance.

On the other hand, only average power of low gamma oscillations in REM sleep was found to be positively correlated with SFI. Since both STN activities in different frequency bands were modulated by sleep stages, we additionally calculated the correlation of SFI and beta, theta and low gamma oscillations in NREM and REM separately (Supplementary Figure 4-5). This generated similar findings: beta and low gamma oscillations in NREM correlated positively with sleep fragmentations severity while only low gamma oscillations in REM demonstrated the same trend (Supplementary Figure 5 a,b,e). This can be attributed to a few reasons which need further investigation. Firstly, REM sleep is not a uniform state and can be divided into phasic and tonic substages. Phasic REM, with enhanced widespread thalamocortical synchronization activities, demonstrated higher arousal index threshold with decreased gamma oscillations than tonic REM. Here, we raised the following hypothesis which need further exploration: the rhythms of phasic and tonic REM can be interrupted in Parkinsonian patients with sleep fragmentation, leading to more tonic REM with lower arousal threshold and higher gamma oscillations. Secondly, previous research found that in chronic sleep deprivation states, enhanced gamma oscillations were modulated by theta activities in compensation for the losses of REM sleep-related synaptic potentiation. Thus, the elevated gamma oscillations in STN LFP may also be a compensatory mechanism of sleep disturbance in Parkinsonian patients.

In summary, our results indicated that the alterations of multiple frequency bands in STN LFPs are involved in sleep disturbance in Parkinsonian patients and combining the low-high power ratio (LHPR) in STN LFPs during sleep may serve as a better biomarker for sleep fragmentation.

Further research would be required to dissect how the oscillatory features in the STN LFPs identified in this study interfered with normal sleep: either through serving as a direct neuropathological biomarker of the neural circuits controlling sleep and wakefulness, or by aggravating nocturnal motor symptoms, cognitive impairment and etc.

### Sleep spindles distinguishable from beta bursts in STN LFPs during sleep

As a hallmark of NREM sleep, sleep spindles have an overlap in frequency band with beta oscillations in normal NREM sleep according to previous researches. A previous study conducted during the perioperative period demonstrated the presence of spindle activity in STN LFPs that synchronized with EEG spindle activityWhat’s more, recent in vivo studies with non-human primates also demonstrated field potential spindles in basal ganglia, which increased during sleep after a learning task especially in the striatum. Here in our analysis, we explored the relationship between beta bursts and spindles recorded in STN LFPs of Parkinsonian patients for the first time, revealing that they are distinguishable in terms of frequency and exhibit a negative correlation in their occupation over time. We confirmed this finding from the following two aspects. First, the peak frequency of beta bursts and sleep spindles in STN were dissociable from each other, with beta bursts showing higher frequency and shorter durations compared with sleep spindles. Second, STN sleep spindles were similar with cortex sleep spindles in peak frequency as well as sleep stage dependent distribution density. Moreover, we contend that it is unlikely to identify STN spindles originating from neighboring pathological oscillation bands, since the spindle activities were absent during wake. However, due to the limited synchronization accuracy of PSG and LFP recording systems, further researches should aim to provide more details about the directionality of information flows between cortical and STN spindles. As sleep spindles have a close relationship with learning and memory, previous studies have found that decreased spindle amplitude and density measured from cortex using EEG were associated with dementia in PD. We further demonstrated a negative correlation between the occurrence of sleep spindles and beta burst in STN LFPs during NREM sleep. This result indicated that besides contributing to sleep fragmentation, beta bursts may also have detrimental effects on spindle related physiological functions such memory consolidation during sleep.

Further research would be required to investigate whether STN spindles, together with above mentioned theta oscillations, could be potential indicators for memory consolidation during sleep and more detailed pathophysiological mechanism of beta oscillations during this process.

### Implications on closed-loop DBS

Closed-loop DBS modulations adjust the stimulation parameters according to the detected biomarkers dynanmically, among which beta power triggered closed-loop DBS in PD have been increasingly studied these days and have been found achieving similar or even better therapeutic effect on motor symptoms comparing with traditional open-loop DBS modulation. However, the effect of beta triggered closed-loop DBS on sleep is unknown and the closed-loop DBS modulation strategy for sleep is less explored. Previous studies have conducted sleep stage classifications based on STN LFPs and provided possibilities towards more precise neuromodulation around sleep-wake cycles. A recent study on the diurnal fluctuations in beta amplitude suggest that the threshold need to be adjusted in beta triggered closed-loop DBS to prevent suboptimal stimulation at night. Here in this study, we showed that with the same threshold defined during awake, beta bursts could still be detected during NREM when patients in Off-DBS and Off-Medication state. Moreover, occurrence of the beta bursts in N2 sleep preceded the transition to stages with more cortical activity (Wake, N1 and REM). These results suggest that, beta power triggered closed-loop DBS, using the same threshold defined during awake, may also reduce sleep fragmentation and improve sleep quality. Another implication of our results on the design of closed-loop DBS is: we may need to differentiate pathological beta oscillation from the sleep spindles. Here we chose beta within the 15-30Hz range to maximize the differentiation from the spindle frequency band. The more commonly used beta range of 13-30Hz were also analyzed to confirm the robustness of our conclusions (Supplementary Figure 6). Further studies would be required to investigate the effect of high frequency STN DBS on sleep spindle and related functions, such as over-night learning and memory consolidation.

### Limitations and Caveats

There are several limitations in our study. First, as all data were acquired in Off-Medication and Off-DBS status, whether these trends were stable under different medication and DBS stimulation states remained unknown. Secondly, as a small-sized clinical study, only 12 patients were included. Thus, further study will include more subjects with different treatment states to further verify the transition patterns in sleep.

## 5. Conclusion

In conclusion, our data revealed features of STN LFP underlying sleep fragmentation in PD. These results deepen our understanding of the mechanism of sleep fragmentations in PD, and offer new insight on how to improve closed-loop DBS in sleep modulation.

## Acknowledgement

We thank Hao Chen, Zongyan Cai, Hao Fang, Yuan Feng and Linze Li for their help with the data recording. We thank Ye Tian and Feng Zhang for their help in the identification of electrodes’ location.

## Contribution

Guokun Zhang: draft of the manuscript, data analysis.

Huiling Yu: conception, draft and edition of the manuscript.

Yue Chen: design of the experiment, data collection.

Chen Gong: data collection.

Hongwei Hao: organization of the research project.

Guo Yi, Shujun Xu, Jianguo Zhang: implementation of surgery.

Yuhuan Zhang, Xuemei Yuan, Guoping Yin: data collection and sleep evaluation.

Huiling Tan: conception, edition and review of the manuscript.

Luming Li: conception, organization of the research project, edition and review of the manuscript.

## Data availability

The original data are not yet openly available, as it is being used in ongoing projects. We welcome enquires for sharing this as part of a collaboration, please contact the corresponding authors.

