## Supplementary for "Neurophysiological Features of STN LFP underlying Sleep Fragmentation in Parkinson’s Disease"

**Supplementary Table 1 Sleep Architecture of Each Participant.**

| Subject | N3 (mins) | N2 (mins) | N1 (mins) | Wake (mins) | REM (mins) | Sleep Efficiency |
|---|---|---|---|---|---|---|
| 1 | 0.5 | 52.5 | 31.5 | 277 | 2 | 0.24 |
| 2 | 15 | 68.5 | 45.5 | 378.7 | 35 | 0.30 |
| 3 | 3 | 226.5 | 60.5 | 164.8 | 67.5 | 0.68 |
| 4 | 58 | 104 | 16 | 209 | 73.5 | 0.55 |
| 5 | 3 | 215 | 76.5 | 189 | 83 | 0.67 |
| 6 | 30.5 | 143 | 33 | 99.5 | 50 | 0.72 |
| 7 | 61 | 123 | 33 | 38.8 | 45.9 | 0.87 |
| 8 | 77.5 | 160 | 20 | 152.6 | 66.5 | 0.68 |
| 9 | 34.5 | 119.5 | 70.5 | 81.5 | 60 | 0.78 |
| 10 | 0 | 172 | 25.5 | 197.4 | 36.1 | 0.54 |
| 11 | 5 | 55.5 | 24.5 | 335.9 | 4.5 | 0.21 |
| 12 | 0 | 138.9 | 42.5 | 40.5 | 119.3 | 0.88 |

**Supplementary Table 2 Sleep Stage Transition Probability.**

| | | To stage … (%) | | | | |
|---|---|---|---|---|---|---|
| | | N3 | N2 | N1 | Wake | REM |
| **Transition from stage …** | N3 | 40.15±38.8 | 42.3±40.16 | 0.29±0.82 | 1.74±3.08 | 0.13±0.48 |
| | N2 | 4.25±4.21 | 84.75±8.61 | 4.99±4.23 | 3.31±3.59 | 2.55±1.71 |
| | N1 | 0.29±0.87 | 28.39±10.86 | 54.59±10.58 | 14.15±6.18 | 1.83±2.17 |
| | Wake | 0 | 0.29±0.75 | 8.85±7.12 | 90.6±7.09 | 0.25±0.42 |
| | REM | 0.05±0.19 | 6.41±8.39 | 3.25±5.97 | 1.83±1.75 | 88.34±11.45 |

**Supplementary Table 3 Sample Size of N2 and REM Transition Process Analysis in Each Participants**

| Subject | N2-N3 | N2-N1 | N2-Wake | N2-REM | REM-N2 | REM-N1 | REM-Wake |
|---|---|---|---|---|---|---|---|
| 1 | 0 | 1 | 4 | 1 | 1 | 0 | 0 |
| 2 | 4 | 2 | 1 | 0 | 2 | 0 | 0 |
| 3 | 3 | 7 | 3 | 4 | 1 | 4 | 3 |
| 4 | 8 | 3 | 0 | 2 | 4 | 0 | 2 |
| 5 | 7 | 7 | 3 | 4 | 0 | 0 | 4 |
| 6 | 5 | 7 | 4 | 2 | 0 | 1 | 1 |
| 7 | 7 | 2 | 0 | 2 | 6 | 0 | 0 |
| 8 | | | | | | | |
| 9 | 2 | 2 | 0 | 4 | 2 | 4 | 3 |
| 10 | 0 | 8 | 5 | 1 | 0 | 2 | 0 |
| 11 | 3 | 3 | 0 | 2 | 0 | 1 | 0 |
| 12 | 0 | 1 | 2 | 5 | 2 | 4 | 1 |

**Supplementary Table 4 p-value of Comparisons for Oscillation Power between Sleep Stages**

| Features | N3-N2 | N3-N1 | N3-Wake | N3-REM | N2-N1 | N2-Wake | N2-REM | N1-Wake | N1-REM | Wake-REM |
|---|---|---|---|---|---|---|---|---|---|---|
| Delta Power | n.s. | *** | *** | *** | *** | *** | ** | n.s. | n.s. | n.s. |
| Theta Power | n.s. | * | *** | n.s. | * | *** | n.s. | * | n.s. | *** |
| Alpha Power | n.s. | n.s. | n.s. | n.s. | * | n.s. | n.s. | n.s. | n.s. | n.s. |
| Beta Power | n.s. | *** | *** | *** | * | *** | * | n.s. | n.s. | n.s. |
| Gamma Power | n.s. | n.s. | *** | *** | * | *** | *** | * | n.s. | n.s. |

*, $p$<0.05; **, $p$<0.01; ***, $p$<0.001; n.s., no statistical significance.

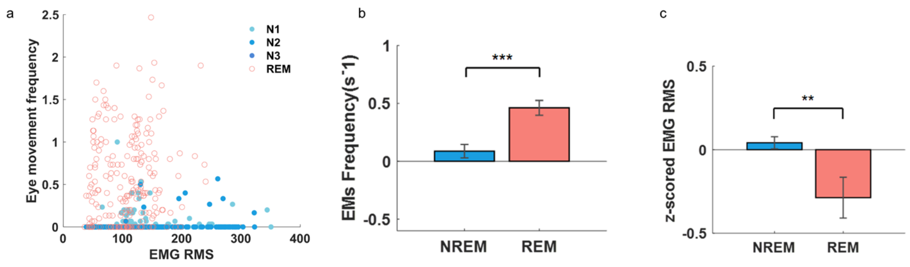


**Supplementary Figure 1.**

**(a)**. EMG RMS and Eye movements (EMs) frequency distribution in one-night PSG; **(b-c)**. Comparison of z-scored EMG RMS and EMs frequency between NREM and RME sleep using data from all subjects' data. The EMs during NREM was significantly less compared to REM sleep, while the EMG RMS was higher during NREM sleep.

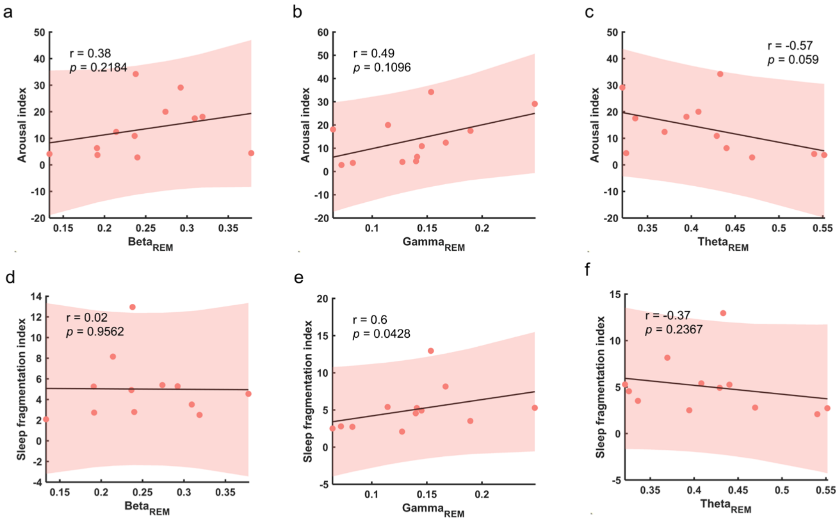


### Supplementary Figure 2

**(a-c)**. Correlations between arousal index through the night and average power of Beta, Gamma and Theta oscillations during REM sleep. Linear fittings and 95% Confidence intervals were shown; **(d-e)**. Correlations between sleep fragmentation index and average power of Beta, Gamma and Theta oscillations during REM sleep. Linear fittings and 95% Confidence intervals were shown.

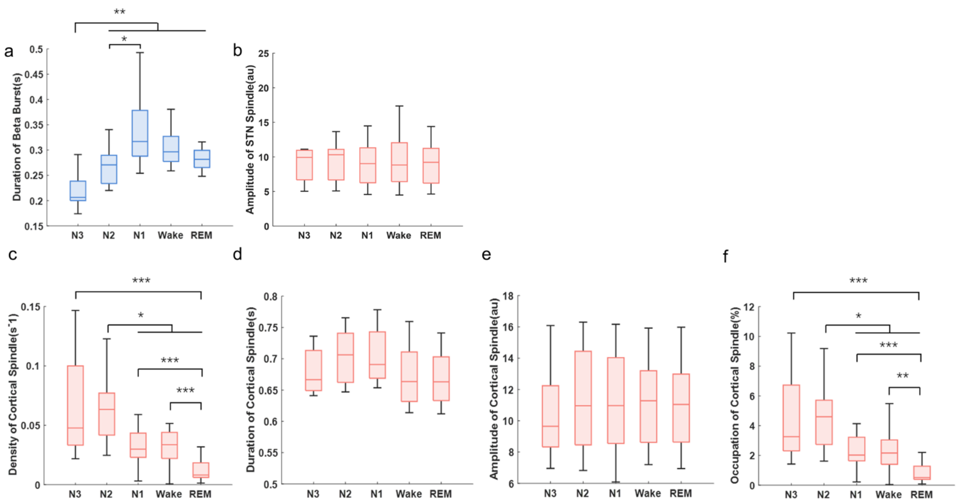


### Supplementary Figure 3

**(a)**. Long Beta bursts amplitude of STN LFP in different sleep stages were averaged within hemispheres and represented as the box-and-whisker plots. The box edges represent the first quartile to the third quartile, with a vertical line drawn through the box at the median. *, $p$<0.05; **, $p$<0.01; **(b)**. Sleep spindles duration of STN LFP in different sleep stages were averaged within hemispheres and represented as the box-and-whisker plots. The box edges represent the first quartile to the third quartile, with a vertical line drawn through the box at the median. No statistical significance was shown; **(c-f)**. Density, duration, amplitude and occupations of cortical spindles during different sleep stages were averaged within hemispheres and represented as the box-and-whisker plots. The box edges represent the first quartile to the third quartile, with a vertical line drawn through the box at the median. *, $p$<0.05; **, $p$<0.01; ***, $p$<0.001.

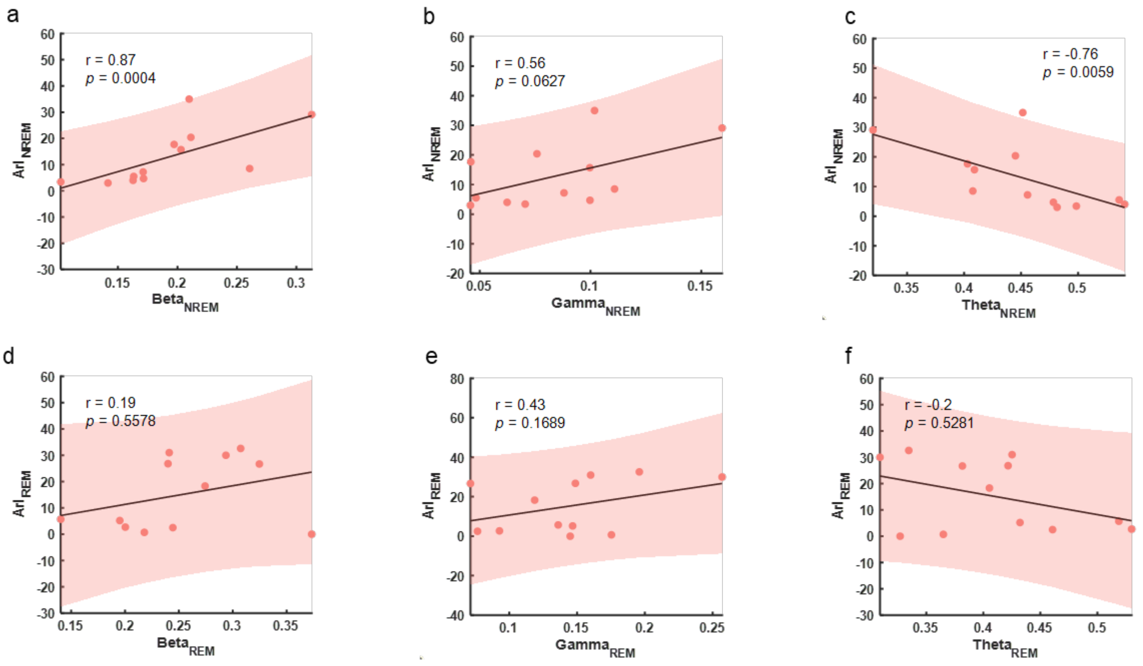


**Supplementary Figure 4**

**(a-c)**. Correlations between arousal index of NREM sleep ($ArI_{NREM}$) and average power of Beta, Gamma and Theta oscillations during NREM sleep; **(d-e)**. Correlations between arousal index of REM ($ArI_{REM}$) sleep and average power of Beta, Gamma and Theta oscillations during REM sleep. Linear fittings and 95% Confidence intervals were shown.

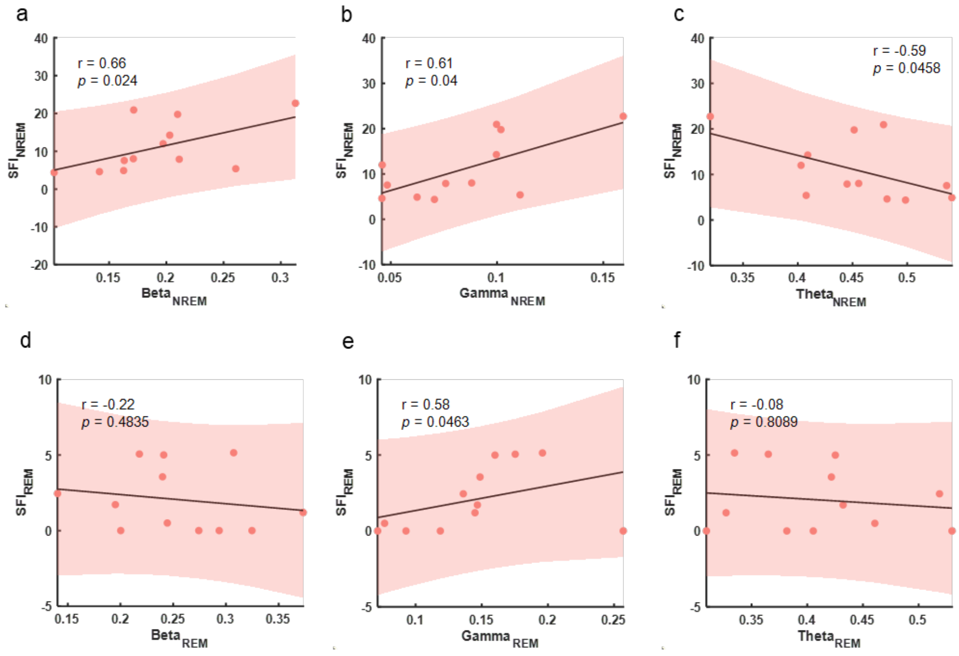


**Supplementary Figure 5**

**(a-c)**. Correlations between sleep fragmentation index of NREM sleep ($SFI_{NREM}$) and average power of Beta, Gamma and Theta oscillations during NREM sleep; **(d-e)**. Correlations between sleep fragmentation index of REM ($SFI_{REM}$) sleep and average power of Beta, Gamma and Theta oscillations during REM sleep. Linear fittings and 95% Confidence intervals were shown.

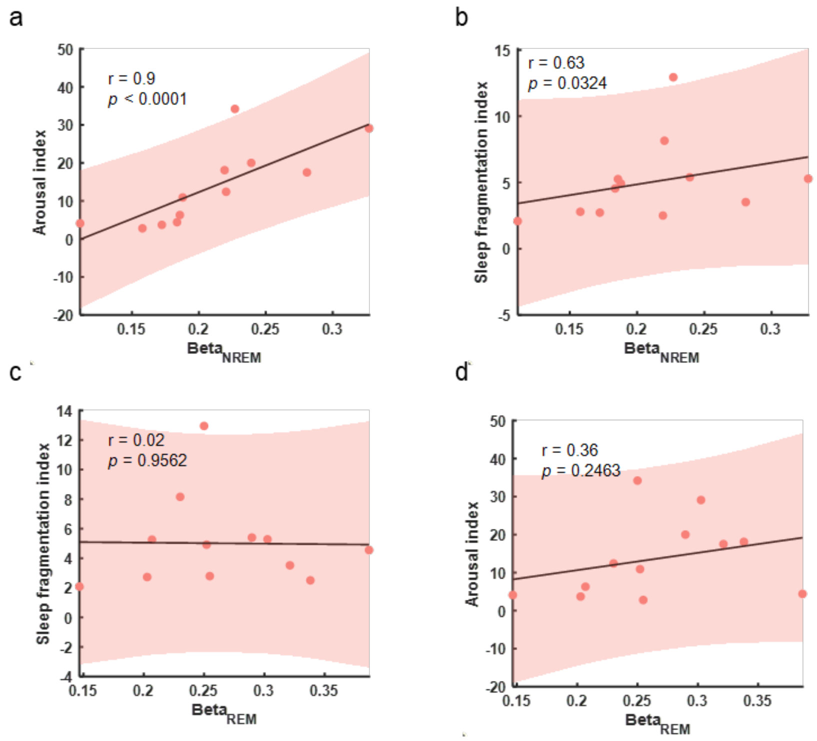


**Supplementary Figure 6**

**(a-b)**. Correlations between average power of beta band during NREM sleep ($Beta_{NREM}$) and Arousal index and sleep fragmentation index through the night respectively. **(c-d)**. Correlations between average power of beta band during REM sleep ($Beta_{REM}$) and Arousal index and sleep fragmentation index through the night respectively. Linear fittings and 95% Confidence intervals were shown.

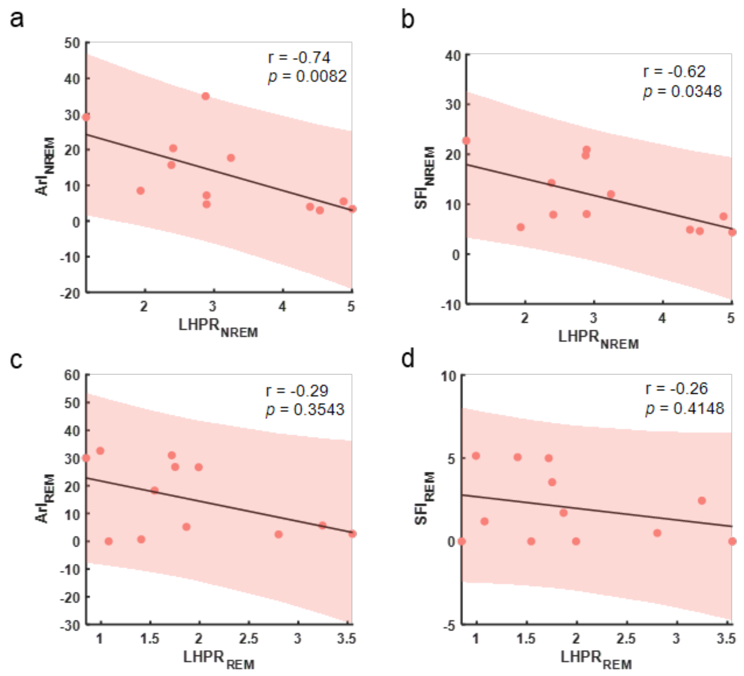


**Supplementary Figure 7**

**(a-b)**. Correlations between LHPR of NREM sleep ($LHPR_{NREM}$) and both arousal index ($ArI_{NREM}$) and sleep fragmentation index ($SFI_{NREM}$) during NREM sleep; **(c-d)**. Correlations between LHPR of REM sleep ($LHPR_{REM}$) and both arousal index ($ArI_{REM}$) and sleep fragmentation index ($SFI_{REM}$) during REM sleep

### Methods

#### *Sleep Evaluation*

The sleep fragmentation was measured by sleep fragmentation index (SFI) and arousal index (ArI). SFI was calculated as the sum of the number of transitions from sleep to wake and transitions from N3, N2, and REM sleep stages to N1, subsequently divided by the total duration of sleep (measured in hours). ArI was calculated as the number of short duration arousals per hour of sleep[1]. Here arousal was defined as a shift in EEG frequency to alpha, theta, and/or frequency >16Hz (but not spindles) lasting at least 3 seconds, with at least 10 seconds of stable sleep preceding the EEG shift. Labeling of arousals during REM requires a concurrent increase in submental EMG lasting at least 1 second[2].

Typical waveforms in the EEG, EMG and EOG measurements that have been previously used for sleep stage labelling in healthy participants can also be observed in recorded PD patients, such as the slow wave activities during N3, K-complex during N2 from EEG channel, as shown in Figure 1c. There was also significant difference in the eye movement frequency and the muscle activities in the submental area, quantified based on the EOG and submental EMG measurements, respectively (Supplementary Figure 1), comparing the REM vs. Non-REM sleep stages. These differences validated the sleep stage labelling.

#### *Signal Preprocessing*

To compare the characteristics of EEG and LFP during sleep stages in parallel, similar preprocessing procedures were applied for both signals. In brief, artifacts were first removed by the following algorithm and then verified manually. For EEG signals, data points with amplitude over a range of ± 120uV were considered muscle and/or eye movement artifacts. For LFP signal, artifacts were detected when the absolute value of the signal amplitude exceeds 10 times the median value. Both EEG and LFPs were bandpass filtered between 0.5Hz-80Hz using an Infinite Impulse response[3,4].

#### *Determination of hemispheres with prominent spindle/beta activity*

This determination was made by identifying the periodic components with peaks within specific frequency ranges (10-17Hz for spindles and 10-35Hz for beta) using FOOOF algorithm[5] with the peak threshold set to be 2, the peak width limits of 2-12, and the aperiodic mode set to be fixed. The peak frequency of spindle and beta was determined as the center frequency of the periodic components that displayed the maximum value relative to the aperiodic components within the aforementioned frequency range.

#### *Spindle Event Detection*

Specifically, the LFP was first bandpass-filtered between 10-17 Hz with a zero-phase Butterworth filter. The amplitude envelope of the filtered signal was then extracted with the Hilbert transform. Candidate spindle events were identified when the envelope amplitude exceeds the threshold of mean+2SD, with the starting and ending range determined by a threshold of mean+0.2SD. To prevent false detections of spindle events caused by an increase in the energy of overall frequency band, candidate spindle events whose envelope amplitudes of 20-30 Hz exceed mean+4.5SD are discarded. Finally, only spindle events with durations between 0.5-2 seconds are retained.

#### *Beta Burst Identification*

LFP signals were decomposed using Morlet wavelet transformation with a 1Hz resolution. The wavelet amplitude within 4Hz range near the beta peak was summed, smoothed (0.2s window) and DC removed. We selected 75% of wavelet amplitude distribution from the LFP signals recorded during daytime wakefulness as the threshold. Segments whose wavelet amplitude surpassed the threshold were considered as beta bursts.
